# A Simple and Robust Cell-Based Assay for the Discovery of Novel Cytokinesis Inhibitors

**DOI:** 10.1101/2020.01.09.900365

**Authors:** Laszlo Radnai, Rebecca F. Stremel, Thomas Vaissiere, William H. Martin, Gavin Rumbaugh, Theodore M. Kamenecka, Patrick R. Griffin, Courtney A. Miller

**Author notes:** Corresponding Author: Courtney A. Miller, Departments of Molecular Medicine and Neuroscience, The Scripps Research Institute, 130 Scripps Way, Jupiter, FL 33458, USA.

## Abstract

Cytokinesis is the last step of mitotic cell division that separates the cytoplasm of dividing cells. Small molecule inhibitors targeting either the elements of the regulatory pathways controlling cytokinesis, or the terminal effectors have been of interest as potential drug candidates for the treatment of various diseases. Here we have developed a cell-based assay for the discovery of novel cytokinesis inhibitors. The assay is performed in a 96-well plate format in 48 hours. Living cells, nuclei and nuclei of dead cells are revealed by a single staining step with three fluorescent dyes, followed by live cell imaging using an automated fluorescence microscope. The primary signal is the nuclei-to-cell ratio (NCR). In the presence of cytokinesis inhibitors, this ratio is expected to increase over time, as the ratio of multinucleated cells increases in the population. Cytotoxicity is quantified as the ratio of dead nuclei to total nuclei. A screening window coefficient (Z’) of 0.65 indicates that the assay is suitable for screening purposes, as the positive and negative controls are well-separated. EC_50_ values for four compounds can be reliably determined in a single 96-well plate by using only six different compound concentrations. An excellent test-retest reliability (R^2^=0.998) was found for EC_50_ values covering a ∼1500-fold range of potencies. The robustness, simplicity and flexibility of the assay is demonstrated here by using modulators of actin dynamics with different mechanisms of action (jasplakinolide, cytochalasin D and swinholide A) and derivatives of the nonmuscle myosin II inhibitor blebbistatin.

## Introduction

In all eukaryotic cells, mitosis is the complex cytological process that separates the already duplicated chromosomes in space into two identical sets, ultimately leading to the formation of two fully functional nuclei in distant parts of the cell^1^. Mitosis is followed by cytokinesis, a process that separates the cytoplasm of dividing cells resulting in the formation of two daughter cells^2, 3^. In animals, a contractile ring of filamentous actin and nonmuscle myosin II (NMII) assembles in the equatorial plane of the dividing cells (between the segregated sister chromatids) that splits the cytoplasm during cytokinesis by forming the so-called cleavage furrow. The process is tightly regulated both in space and time in a cell-type specific manner^2-5^. Cytokinesis inhibitors targeting actin, myosin or other regulatory and structural elements necessary for the proper functioning of the cleavage furrow have not only provided useful information about the function of these elements, but are increasingly recognized as potential drug candidates^6^.

Here we present a cell-based assay that is amenable to the discovery of novel cytokinesis inhibitors. The assay is run in 96-well plates with a total incubation time of 48 hours. We demonstrate the robustness, flexibility and reliability of the assay by using small molecule inhibitors of NMII (blebbistatin, *para*-aminoblebbistatin and *para*-nitroblebbistatin) and modulators of actin polymerization with different mechanisms of action (jasplakinolide, cytochalasin D and swinholide A). Blebbistatin is an inhibitor of NMII that blocks cellular blebbing and disrupts directed cell migration^7^. It also blocks cytokinesis by inhibiting the contraction of the cleavage furrow. This results in the accumulation of binucleated cells in the population^7^. *Para*-nitroblebbistatin and *para*-aminoblebbistatin are photostable derivatives of blebbistatin with reduced cytotoxicity^8, 9^. *Para*-aminoblebbistatin also shows highly improved water solubility^9^. Jasplakinolide is a stabilizer of pre-formed actin filaments and an inducer of actin polymerization *in vitro*^10, 11^. By facilitating filament nucleation and lowering the critical actin concentration, jasplakinolide treatment results in the formation of amorphous actin masses in cultured cells^11^. Cytochalasin D inhibits actin polymerization *in vitro* by binding to the net polymerizing ends of actin filaments and blocking the addition of new monomers to these sites^12, 13^. It also binds actin monomers, inducing the formation of dimers and thereby increasing the rate, but decreasing the extent of polymerization^14, 15^. In cells, cytochalasin D treatment leads to the disruption of the cytoskeletal actin filament network and the aggregation of actin filaments^16, 17^. The third modulator of actin polymerization, Swinholide A, is a highly potent toxin that rapidly severs actin filaments *in vitro*^18^, and disrupts the actin cytoskeleton of living cells *in* vivo^18, 19^.

## Materials and Methods

### Cell cultures

Frozen aliquots of COS7 cells (cat. CRL-1651, American Type Culture Collection, Manassas, VA) were thawed and immediately diluted ten-fold in culture medium containing 89% Dulbecco’s Modified Eagle Medium, 10% Fetal Bovine Serum, and 1% Antibiotic-Antimycotic solution (cat. 11995073, cat. 26140079, and cat. 15240062, Life Technologies Carlsbad, CA). The cell suspension was centrifuged at full speed (7,197 × g) at 20 °C for 10 minutes in a refrigerated centrifuge (cat. 5430 R, Eppendorf, Hauppauge, NY). The supernatant was discarded, and the pellet was resuspended in culture medium at a final density of ∼50,000 cells/ml. Cells were plated onto 75 cm^2^ flasks (cat. 430641U, Corning, Corning, NY) at a density of 500,000 cells/flask. Following 3 days of incubation (37 °C and 5% CO_2_), old media was removed from the flasks and the cell layers were washed twice with 5 ml of Dulbecco’s phosphate-buffered saline (cat. 14190250, Life Technologies, Carlsbad, CA). Subsequently, 2 ml of Trypsin-EDTA (0.25%) solution (cat. 25200072, Life Technologies, Carlsbad, CA) was added to each flask to dissociate cells. Flasks were incubated at 37 °C for 10 minutes to allow detachment of cells from the surface. Cell dissociation was further facilitated by pipetting the suspension up and down several times until no cell aggregates were observed by visual inspection under a stereomicroscope. To inhibit trypsin, 8 ml of fresh culture medium was combined with 2 ml of cell suspension. Cell density was determined by counting the cells in a hemocytometer. The suspension was diluted to a density of 20,000 cells/ml. Cells were immediately plated onto flat bottom, 96-well cell culture plates (cat. 25-109, Genesee Scientific, El Cajon, CA) by transferring 100 μl of suspension to each well using a multichannel pipette resulting in a final surface density of 2,000 cells/well.

### Treating cells

Following 24 hours of incubation (37 °C, 5% CO_2_), cells were treated with jasplakinolide (cat. 2792/100U, R&D Systems (Minneapolis, MN), cytochalasin D (cat. 11330-5, Cayman Chemical, Ann Arbor, MI), swinholide A (cat. 501146229, Fisher Scientific, Hanover Park, IL), blebbistatin, *para*-aminoblebbistatin or *para*-nitroblebbistatin prepared in compound plates (described below). Blebbistatin derivatives were provided by Albany Molecular Research Inc., Albany, NY (custom synthesis). First, compounds of interest were dissolved in dimethyl sulfoxide (DMSO, cat. D2650, Sigma-Aldrich, St. Louis, MO). Second, six-step serial 1:2 dilutions of compound solutions were prepared in DMSO using solvent-resistant polypropylene microplates (cat. 3357, Corning, Corning, NY). Concentration ranges (125-3.91 nM, 500-15.63 nM, 10-0.31 nM, 10-0.31 μM, 10-0.31 μM, and 10-0.31 μM for jasplakinolide, cytochalasin D, swinholide A, blebbistatin, *para*-aminoblebbistatin and *para*-nitroblebbistatin, respectively) were determined based on preliminary results. A 2 mM solution of *para*-aminoblebbistatin in DMSO was also prepared and used as positive control, while pure DMSO was used as a negative control. Third, compound plates were prepared by transferring 2.4 μl of positive and negative controls and compound solutions into each well of a 96-well plate containing 117.6 μl of culture media (50-fold dilution) using a multichannel pipette. Wells in the first and last rows were used exclusively for negative and positive controls. Solutions were mixed by shaking the compound plate for 1 min at room temperature at 1200 rpm using a microplate shaker (cat. 12620-926, VWR, West Chester, PA). A typical plate layout map used for screening experiments is shown in Figure 1. Finally, 100 μl of diluted compound solutions were transferred from the compound plate to the assay plate (containing cell cultures in 100 μl of culture media) using a multichannel pipette (2-fold dilution). NOTE: The DMSO concentration in the compound and assay plate was 2% and 1%, respectively. The final concentration of the positive control *para*-aminoblebbistatin was 20 μM, resulting in ∼100% signal. (See Fig. 2. and Results for more details.) All measurements were carried out in triplicate.

**Figure 1.**
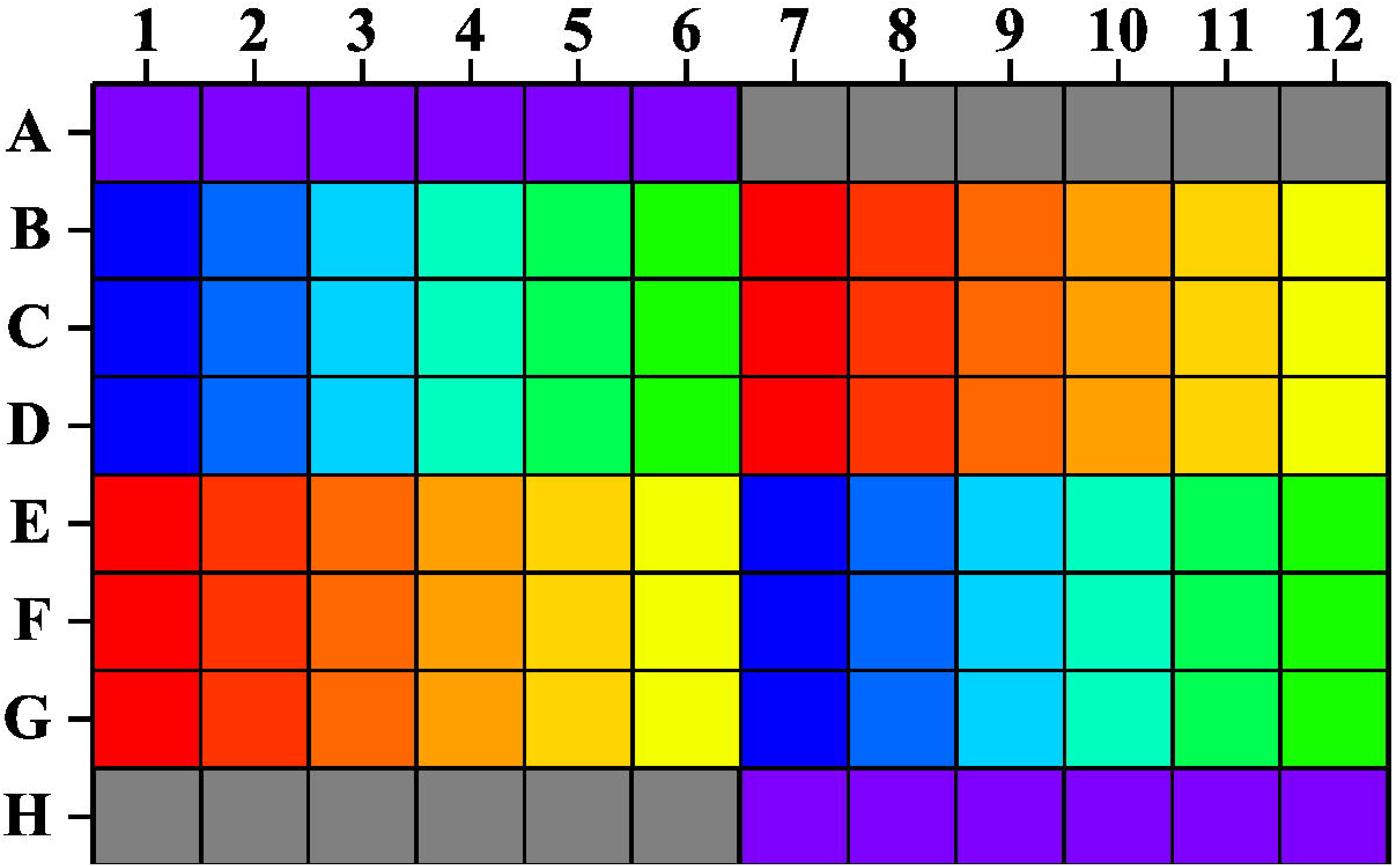
Assay plate layout used in screening experiments. Row A and row H were reserved for negative (1% DMSO) and positive (20 μM *para*-aminoblebbistatin, 1% DMSO) controls (violet and gray, respectively). Six-step serial 1:2 dilutions of four different compounds were prepared in DMSO and diluted in culture media. Subsequently, diluted compound solutions were transferred to the assay plate, such that every experiment was performed in triplicate.Blue-to-green and red-to-yellow color gradients (wells 1-6 and 7-12, rows B-D and E-F) represent the applied concentration gradients.

**Figure 2.**
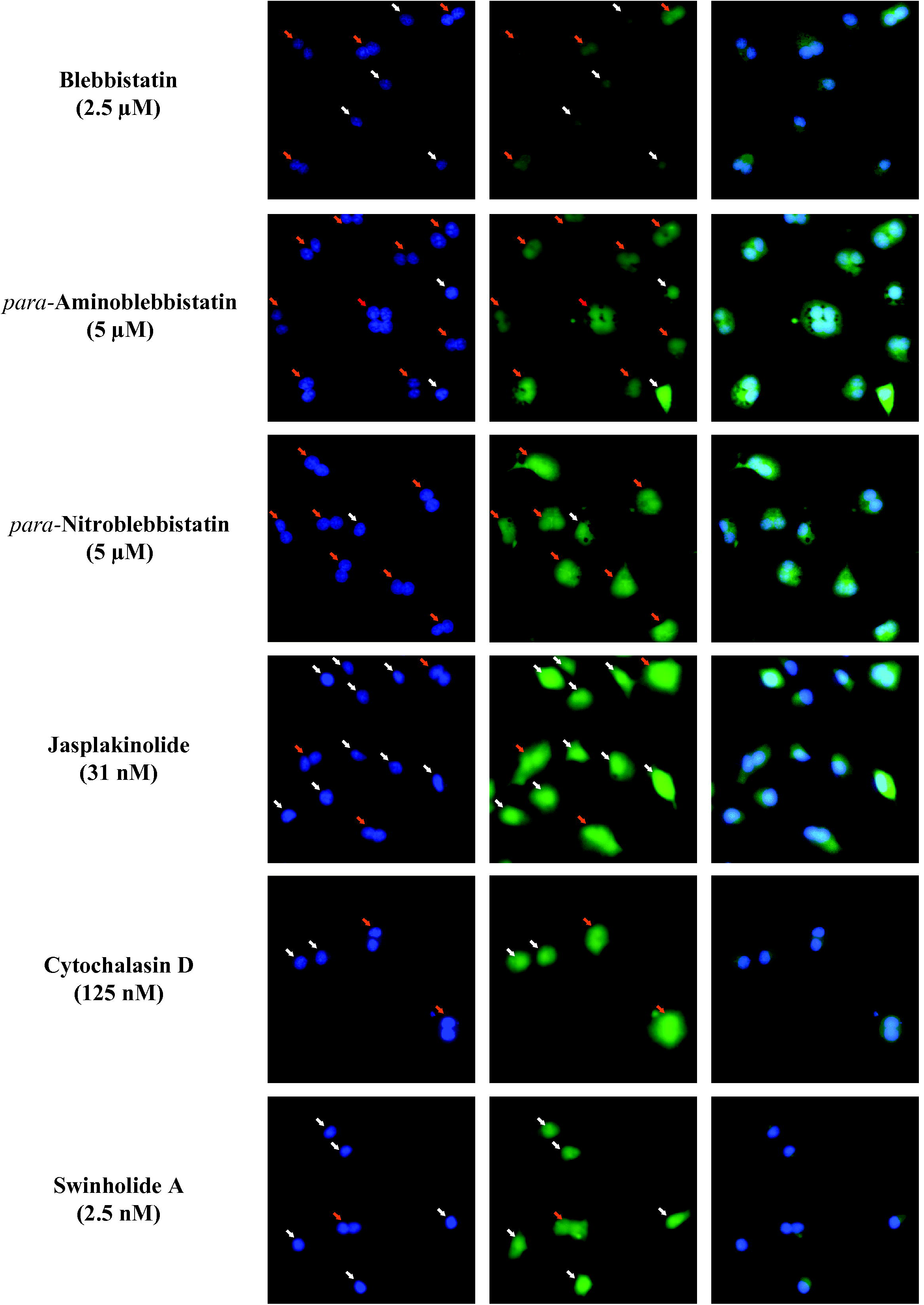
(A) Representative images of *para*-aminoblebbistatin (20 μM) treated and control cells showing living cells (green, FDA staining), nuclei (blue, Hoescht33342 staining), and nuclei of dead cells (red, PI staining), with overlaid images shown in the far-right panels. Most of the *para*-aminoblebbistatin treated cells are bi- or multinucleated and much bigger than negative control cells, which are primarily mononucleated. Examples of a mono- (white arrows) and a binucleated (orange arrows) cell are highlighted in each image. Far more binucleated cells are present in the *para*-aminoblebbistatin treated condition. Note that there is no living cell body revealed around the nuclei showing PI staining (white circles). However, these nuclei are positive for both Hoescht33342 and PI, confirming their identity. Images were not manipulated other than changing the brightness and contrast for visibility. (B) Potential artifacts may result from compound precipitation and object misidentification. COS7 cells were treated with *para*-nitroblebbistatin and blebbistatin at 30 μM. Left and right panels show raw images and detected objects (nuclei – yellow, living cell – red, dead nuclei – cyan), respectively. All three channels are shown for both compounds. Crystals of *para*-nitroblebbistatin showed signal in the green and red channels (yellow circles), while blebbistatin crystals showed blue fluorescence (cyan circles). Accordingly, the *para*-nitroblebbistatin crystal cluster shown was initially misidentified as a living cell and also as a dead nucleus, while the blebbistatin crystal was misidentified as an elongated nucleus. Based on cell shape and size, parts of the cytoplasm can also be misidentified as separate cells (orange circles). However, since those “cells” not overlapping with nuclei, those “nuclei” not overlapping with cells or those “dead nuclei” not showing labeling for both Hoechst33342 and PI are excluded from our calculations, these artifacts usually have only limited effect on our signals. (C) *Para*-aminoblebbistatin was selected as a positive control due to its high solubility, photostability and low cytotoxicity^9^. By fitting the dose response data to the Hill-equation, an EC_50_ of 5.3 μM with a very steep transition (Hill-constant = 7.7) was determined. The signal approaches its maximum value around and above ∼10 μM compound concentration, therefore, 20 μM *para*-aminoblebbistatin was chosen as a reliable positive control in this work. (The red horizontal line represents the cytotoxicity threshold of 0.015. See text for further explanation.) (D) A Z’^26^ of 0.65 was determined for a half plate of negative (1% DMSO) and half plate of positive controls (1% DMSO, 20 μM *para*-aminoblebbistatin), indicating a reliable assay for screening with well-separated positive and negative controls.

### Staining cells

Following 24 hours of incubation in the presence of compounds (37 °C, 5% CO_2_), cells were stained by fluorescein diacetate (FDA, cat. F7378, Sigma-Aldrich, St. Louis, MO), Hoescht33342 (cat. H3570, Life Technologies, Carlsbad, CA) and propidium iodide (PI, cat. P3566, Life Technologies, Carlsbad, CA) in a single step. Staining solution was prepared by combining 9 μl FDA (24 mM stock solution in DMSO), 22.2 μl Hoescht33342 (16.2 mM stock solution in water), 96 μl PI (1.5 mM stock solution in water) and 11.873 ml culture medium in a 50 ml centrifuge tube for each plate. To each well of the assay plate containing 200 μl of treated cell culture, 100 μl of staining solution was added using a multichannel pipette. This resulted in a final concentration of 6 μM, 10 μM, and 4 μM for FDA, Hoescht33342 and PI, respectively. Assay plates were incubated for 10 minutes (37 °C, 5% CO_2_). Finally, the staining solution was replaced by fresh culture medium in each well (100 μl/well).

### Imaging

An IN Cell Analyzer 6000 automated fluorescence microscope (cat. 29-0433-23, GE Healthcare Bio-Sciences, Marlborough, MA) was used for imaging cells. Since the binucleated phenotype remained detectable without any observable change in cell morphology or the number of nuclei for hours at room temperature, cells were not incubated during imaging. Image acquisition was performed using the Nikon 10X/ 0.45, Plan Apo, CFI/60 objective. Lasers operating at 405 nm, 488 nm, and 561 nm in conjunction with 455 nm, 525 nm and 605 nm emission filters, were used for the visualization of Hoescht33342, fluorescein and PI signals, respectively. Cells were counted while imaging to make sure that at least 600 cells were imaged per well. Lower cell counts may yield unreliable results. (See Results and Discussion for a detailed statistical explanation.)

### Data analysis

Images were analyzed automatically by the IN Cell Developer Toolbox software (cat. 25-8098-26, GE Healthcare Bio-Sciences, Marlborough, MA). Nuclei of living and dead cells were identified by using the corresponding fluorescent signal of Hoescht33342. Nuclear segmentation was performed using the following parameters: minimum target area: 85 μm^2^, sensitivity: 50, sensitivity range 1.30, precise mask: enabled. Steps of postprocessing: watershed clump breaking, erosion (kernel size 9), sieve (keep targets with an area greater than 20 μm^2^), border object removal. Cell bodies of living cells were identified by using the corresponding fluorescent signal of fluorescein. Cytoplasm segmentation was performed using the following parameters: noise suppression: heavy, remove shading artifacts with area greater than: 2960 μm^2^, use octagonal morphology: enabled. Steps of postprocessing: sieve (keep targets with an area greater than 200 μm^2^), erosion (kernel size 3), watershed clump breaking, fill holes, dilation (kernel size 3), sieve (keep targets with an area greater than 200 μm^2^), border object removal. Nuclei of dead cells were identified by using the corresponding fluorescent signal of PI. Nuclear segmentation was performed using the following parameters: minimum target area 52 μm^2^, sensitivity: 20, sensitivity range 1.30, precise mask -enabled, use octagonal morphology: enabled. Steps of postprocessing: watershed clump breaking, erosion (kernel size 5), sieve (keep targets with an area greater than 20 μm^2^), border object removal.

A cytokinesis inhibitor allows nuclei to divide, but blocks the separation of cell bodies, resulting in the formation of bi- and multinucleated cells. The inhibitory effect of compounds can be estimated and compared by calculating the ratio of nuclei to cell numbers in the living cell population. Each living cell must contain at least one nucleus and each nucleus must belong to a living cell in this population. Therefore, “cells” not overlapping with nuclei (potentially misidentified cells) and “nuclei” not overlapping with living cells (dead and potentially misidentified nuclei) were excluded from the calculation of the nuclei-to-cell ratio (NCR). This step was crucial to avoid several artifacts (see Results and Discussion). In addition to cytokinesis inhibition, compounds may show cytotoxic effects, which can affect the primary signal (e.g. through nuclear fragmentation or selective death of multinucleated cells). Therefore, cytotoxicity was also quantified as the ratio of dead nuclei to total nuclei. Dead nuclei must show positive staining for both Hoescht33342 and PI. Therefore, all those objects not showing double labeling were excluded from this calculation. Similarly, the total number of nuclei was calculated as the sum of Hoechst33342 positive nuclei overlapping with cells or overlapping with PI positive nuclei. OriginPro 2017 software (cat. ORGNPRO, OriginLab, Northampton, MA) was used to plot the NCR against the compound concentration and to determine the half maximal effective concentration (EC_50_) by fitting the 6-point dose-response data to the Hill-equation:

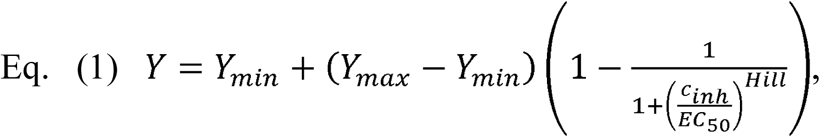

where *Y* is the NCR, *c*_*inh*_ is the concentration of the compound, *Y*_*min*_ is the NCR in the absence of inhibitors, *Y*_*max*_ is the extrapolated value of the NCR at 100% inhibition, and *Hill* is the Hill-constant.

## Results and Discussion

The presence of cytokinesis inhibitors in a cell culture leads to the accumulation of bi- and multinucleated cells. For example, treating COS7 cells with 20 μM *para*-aminoblebbistatin for 24 hours resulted in a cell population where most cells were binucleated (Fig. 2A). To visualize the cytoplasm, we used FDA, a membrane-permeant non-fluorescent cell viability dye that is hydrolyzed to fluorescein in living cells^20^ by naturally present esterase enzymes^21^. The resulting product fluorescein accumulates in the cytoplasm where it shows a bright green fluorescence^20^, thereby providing an excellent signal-to-noise ratio. Nuclei were visualized by Hoescht33342, a cell-permeable DNA stain^22^. The excitation and emission peaks of Hoescht33342 (ex: 350 nm, em: 461 nm, DNA-bound) and fluorescein (ex: 490 nm, em: 526 nm) are well separated from each other. Therefore, these dyes can be used in parallel in imaging applications. We realized that not all nuclei revealed by Hoescht33342 staining overlapped with living cells (see the highlighted areas in Fig. 2A). To confirm the identity of these objects, we used a second DNA dye, PI^23^, which is membrane-impermeant^24^. This dye is widely used to stain nuclei of dead cells in a cell population^25^. The spectral properties of PI (ex: 535 nm, em: 617 nm, DNA-bound) enabled us to use it in parallel with our other two dyes.

To quantify the inhibitory effect of our compounds, we used the IN Cell Developer Toolbox software (GE Healthcare Bio-Sciences) to automatically count nuclei, living cells, and dead nuclei (Hoescht33342-, fluorescein-, and PI-positive objects, respectively). Then, we calculated the nuclei-to-cell ratio (NCR) for each sample (cells from the same well) by simply dividing the number of those nuclei overlapping with cells (living nuclei) with the number of those cells overlapping with nuclei. (Every cell contains at least one nucleus and every nucleus belongs to a cell.) The NCR is expected to be greater than one for any cell cultures (even for untreated ones) because there is always a subpopulation of dividing cells containing two nuclei. Cytotoxicity was also quantified as the ratio of dead nuclei to all nuclei. Here, only the double-labeled (Hoechst33342 and PI positive) nuclei were counted as dead nuclei. The total number of nuclei was calculated as the sum of the double-labeled dead nuclei and the above defined living nuclei. With this strategy, several artifacts resulting from object misidentification could be efficiently avoided.

One main source of such artifacts is the limited solubility of the compounds in culture media. This is generally unknown in screening projects. Aqueous solubility (if available) may not match the solubility in culture media at 37°C. Applying compounds at concentrations higher than the solubility may result in compound precipitation. Precipitates may show fluorescence in one or more channels and may be similar in shape and size to cells or nuclei. Therefore, they are prone to be misidentified as such. To demonstrate the effect of compound precipitation, we treated COS7 cells with *para*-nitroblebbistatin and blebbistatin at 30 μM. The kinetic aqueous solubility of blebbistatin and *para*-nitroblebbistatin has been reported to be 9.3 μM and 3.6 μM, respectively^9^. Since 30 μM was much higher than the solubility in both cases, compounds precipitated, and fluorescent crystals attached to the surface were detected (Fig. 2B). Crystals of *para*-nitroblebbistatin showed bright green fluorescence and also detectable signal in the red channel. Blebbistatin crystals showed bright blue fluorescence. Accordingly, *para*-nitroblebbistatin crystals were prone to be misidentified as living cells and also as dead nuclei. Blebbistatin crystals were prone to be misidentified as nuclei. However, our method was largely resistant to these artifacts since we used double fluorescent labeling for every object of interest (living cells must overlap with nuclei; nuclei must overlap with either living cells or dead nuclei; dead nuclei must overlap with nuclei). This strategy also helped to avoid misidentification of any parts of the cytoplasm as a separate cell (Fig. 2B) or misidentification of any potential fluorescent contamination present in the field of view.

Even for cell populations where no precipitation-related artifacts were present, the number of misidentified objects was effectively reduced by applying the above-mentioned rules. For example, ∼90,000 and ∼110,000 Hoechst33342-positive objects, ∼68,000 and ∼62,000 fluorescein-positive objects, and ∼1,000 and ∼1,300 PI-positive objects were identified for 96 wells of negative (1% DMSO) and positive controls (1% DMSO, 20 μM *para*-aminoblebbistatin; see below) across 8 plates, respectively. To calculate NCR, only ∼94% and ∼93% of Hoechst33342-positive objects (“nuclei”, overlapping with fluorescein-positive objects) and 97% and 88% of fluorescein-positive objects (“living cells”, overlapping with Hoechst33342-positive objects) were used, respectively. To calculate cytotoxicity, only ∼60% and 57% of PI-positive objects (“dead nuclei”, PI and Hoechst33342 double-positive), and ∼95% and ∼94% of Hoechst33342-positive objects (“total number of nuclei”, Hoechst33342 positive objects overlapping either with fluorescein- or PI-positive objects) were used, respectively. The rejected objects were either misidentified fluorescent objects, or correctly identified parts of cell bodies or nuclei along the edges of the image where the overlapping nuclei or cell bodies had been rejected during image analysis. (It is impossible to correctly assign the phenotype if the whole cell is not visible.)

Precipitation may not only lead to fluorescent artifacts, but also limit the effective concentration of the compound, resulting in limited signal in single-point or dose-response experiments. Moreover, precipitates are usually not tolerated well by living cells. Elevated cytotoxicity may be a good indicator of this problem. The intrinsic toxicity of the compound might also result in higher levels of observed cytotoxicity. Significant levels of cytotoxicity can only be determined empirically. Negative and positive controls across 8 plates (96 wells each) showed an average cytotoxicity of 0.007±0.009 and 0.007±0.005, respectively. More than 90% of the data points were under the level of 0.015 in both categories. Therefore, a cytotoxicity signal equal to or greater than ∼0.015, corresponding to ∼1.5% dead nuclei in the population, was considered significant. Around and above this threshold, deterioration of the NCR signal was usually observed (see Fig. 4. D and F for examples), further supporting the idea that our threshold was set to the appropriate level. Besides the limited effective concentration due to solubility issues, selective loss of multinucleated cells, cell clustering, cell detachment from the surface, abnormal morphology or nuclear fragmentation may cause deviation of the NCR signal from the expected value. Analyzing any signal where the associated cytotoxicity is too high should be generally avoided.

*Para*-aminoblebbistatin is known as a non-cytotoxic derivative of the myosin inhibitor blebbistatin, showing greatly improved photostability and solubility^9^, making it optimal as a positive control in our assays. The potency (EC_50_) of *para*-aminoblebbistatin was determined in dose-response experiments (Fig. 2C). An EC_50_ of 5.3 μM was found by fitting the NCR signal to the Hill equation (Eq. 1). Previously, *para*-aminoblebbistatin was shown to inhibit the proliferation of HeLa cells with a similar EC_50_ value (∼18 μM)^9^. Interestingly, the dose-response curve determined here showed a very steep transition (Hill constant = 7.7), such that the signal approached 100% around 10 μM compound concentration. To ensure that the inhibition level is always around 100%, we chose 20 μM *para*-aminoblebbistatin as a positive control. Consistent with the literature^9^, no significant cytotoxicity was observed up to 200 μM concentration. The Z’ (also known as the screening window coefficient) is a widely used statistical parameter that helps to assess the reliability of screening data^26^. We determined the Z’ for a half plate of negative (1% DMSO) and a half plate of positive (1% DMSO, 20 μM *para*-aminoblebbistatin) controls:

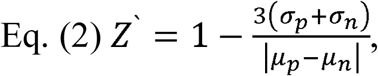

where σ_n_, σ_p_ and μ_n_, μ_p_ are the sample standard deviations and means of the negative and positive controls, respectively (Fig. 2D). We found a Z’>0.5 (Z’=0.65), which indicates that the assay is excellent for screening purposes^26^.

Next, we treated COS7 cells with dilution series of blebbistatin, *para*-aminoblebbistatin, *para*-nitroblebbistatin, jasplakinolide, cytochalasin D or swinholide A. Although these compounds have different molecular targets and mechanisms of action, they all inhibit cytokinesis^7-9, 18, 27, 28^. Accordingly, cells developed similar, bi- and multinucleated phenotypes in response to all compounds (Fig. 3). Although slightly different cellular morphology was observed, our protocol was effective in the quantitation of the multinucleated phenotype in all cases.

**Figure 3.**
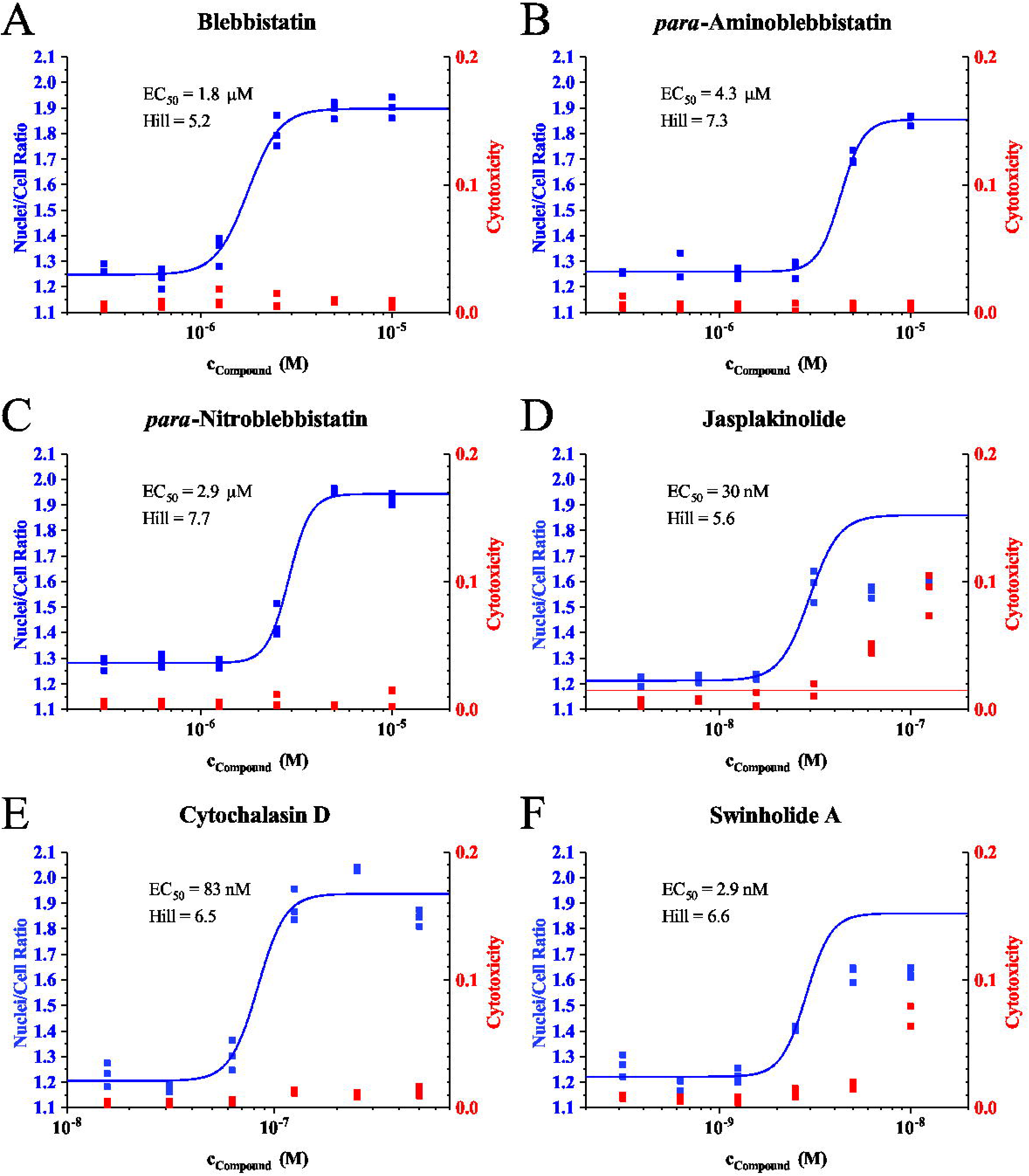
COS7 cells treated with NMII inhibitors (blebbistatin, *para*-aminoblebbistatin, or *para*-nitroblebbistatin) or actin polymerization modulators (jasplakinolide, cytochalasin D, or swinholide A) developed similar multinucleated phenotypes. Compounds were applied at the concentrations shown in parentheses and the cells were photographed after 24 hours of incubation. Images of nuclei (blue, left panels) and living cells (green, middle panels) were chosen to include both mono- (white arrows) and binucleated (orange arrows) cells. Occasionally, cells with more than two nuclei were observed in these experiments (red arrows). Overlaid images are shown to guide the eyes (right panels). Images were not manipulated other than changing the brightness and contrast for better visibility. PI-positive (dead) nuclei were not observed in these fields of view.

Dose-response curves with very steep transitions (Hill-constants between 5.2 and 7.7) were observed (Fig. 4). By using only six-step serial 1:2 dilutions and performing every experiment in triplicate, it was possible to obtain reliable EC_50_ data for four compounds on a single 96-well plate (For EC_50_ values, see Fig. 4. and the upper right panel in Fig. 5. See Fig. 1 for plate layout.). Repeated experiments with fresh cell cultures yielded similar EC_50_ values (Fig. 5). To quantitate the reliability of the assay, linear regression analysis was performed using the data from both runs. A coefficient of determination of 0.998 showed excellent test-retest reliability (Fig. 5).

**Figure 4.**
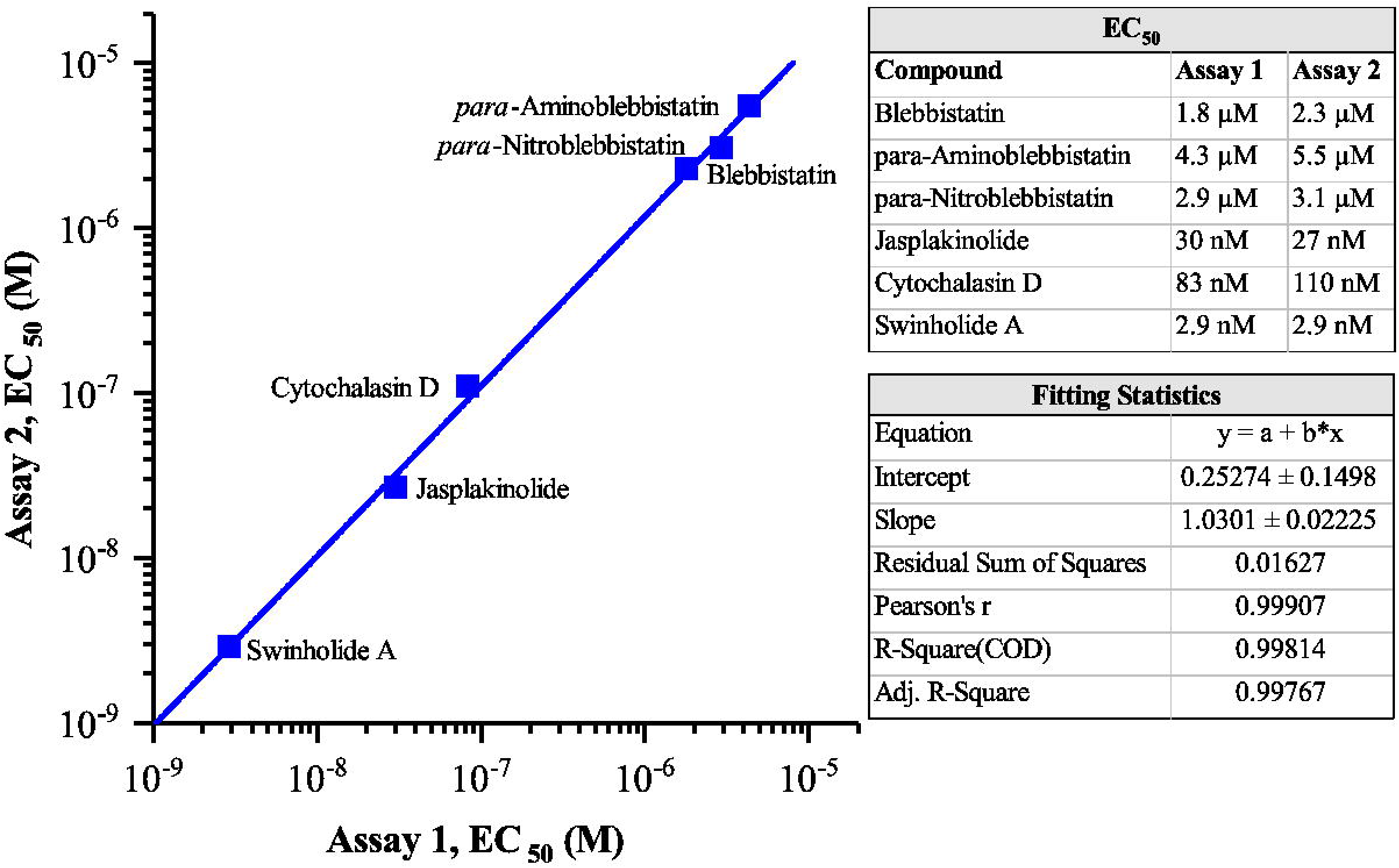
Representative dose-response curves of blebbistatin (A), *para*-aminoblebbistatin (B), *para*-nitroblebbistatin (C), jasplakinolide (D), cytochalasin D (E) and swinholide A (F). Only cells overlapping with nuclei and nuclei overlapping with cells were counted to calculate the primary signal (NCR). Cytotoxicity is calculated as the ratio of dead nuclei to total nuclei. (Here, “total nuclei” is the sum of Hoescht33342 and PI double-positive nuclei and Hoescht33342-positive nuclei overlapping with living cells.) The primary signal was analyzed by fitting the experimental data to the Hill-equation. Data points where the associated cytotoxicity signal was above the empirical threshold of 0.015 (red horizontal line, representing a dead nuclei ratio of 1.5%) were disregarded during the fitting process. This threshold usually represents levels of cell death that affects the primary signal (see 10 and 5 nM swinholide A, or 125 and 62.5 nM jasplakinolide as examples). In these cases, it was necessary to assume that the Y_max_ parameter is equal to the signal of positive control and keep it fixed during the fitting process. In some cases, it might also be necessary to fix the Hill constant to get a reasonable fit.

**Figure 5.**
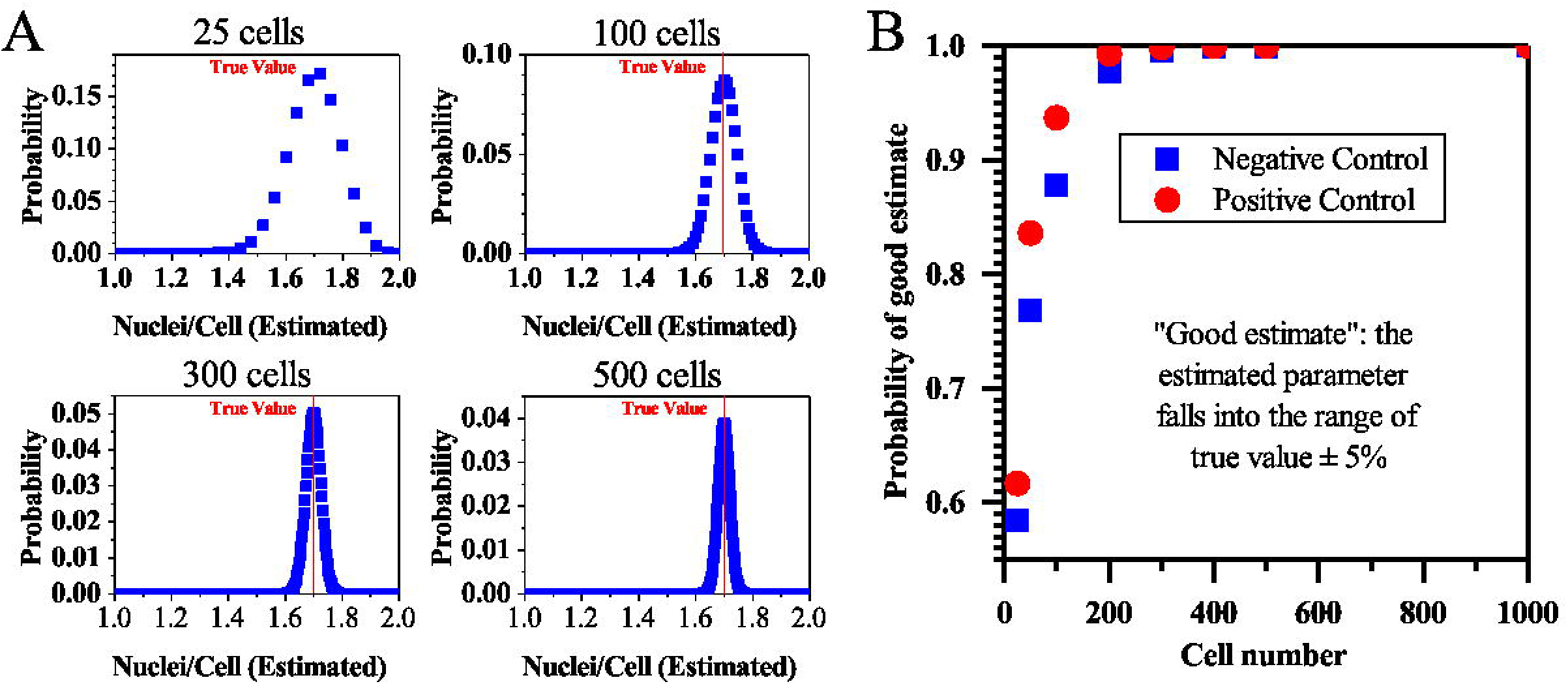
Test-retest reliability of the cytokinesis assay. EC_50_ values (upper right panel) for blebbistatin, *para*-aminoblebbistatin, *para*-nitroblebbistatin, jasplakinolide, cytochalasin D, and swinholide A were determined in two independent runs (Assay 1 and Assay 2). Compounds showed potencies in a ∼1500-fold range. Linear regression analysis (lower right panel) revealed a coefficient of determination (“R-square”) of 0.998 showing that the assay has excellent reliability.

Note that the “dense sampling” of the dose response curves (1:2 dilutions), together with the observed steep transitions constrain the possible values of the EC_50_ in a less than two-fold range, even if other parameters are not well constrained. For example, fixing the Hill-constant at any value from ∼5 to arbitrarily high numbers usually yields fits with similar goodness, while only slightly affecting the determined EC_50_. This does not represent a problem if the goal is to estimate the EC_50_ accurately. To improve the accuracy of Hill-constant estimation, one would need more data points in the transition zone of the dose response curve, at the cost of fewer compounds being tested in parallel on the same plate. It may also be necessary to fix the *Y*_*max*_ to get reasonable fits. Such situations are shown in Fig. 4E and Fig 4F, where cytotoxicity resulted in signal deterioration and limited the useful data available for fitting. Therefore, the *Y*_*max*_ was made equal to the average NCR of the positive control and kept fixed during the fitting process. This strategy assumes that the cell division proceeds at similar rates in all these experiments, which seems to be valid based on the similar maximal NCR ratios observed for blebbistatin derivatives and cytochalasin D, but it may not be universally valid. Even if the validity is questionable, the dense sampling of the dose response curves makes it very unlikely to introduce more than two-fold uncertainty around the EC_50_ values.

There are several critical steps in our protocol. First, finding the optimal cell density is crucial. Low cell density may lead to slow cell growth, unnecessarily long incubation times and insufficient total numbers of cells to estimate the signal accurately (see statistical considerations below). On the other hand, high cell density makes it difficult for any automated algorithm to correctly identify cells due to extensive overlap. This usually leads to an apparent increase of the NCR, as multiple cells are recognized as one cell. Therefore, optimal cell density must be determined in separate experiments for every particular cell line used. For COS7 cells, we found that a surface density of 2,000 cells/well on 96 well plates gave optimal results.

Second, it is highly recommended that compound solutions be diluted in media before adding them to the cell cultures. This allows the solutions to be mixed thoroughly by shaking the plate, until a homogenous mixture that is relatively dilute for DMSO (2% in this work) forms. This can safely be applied to the cell cultures, as the relatively large, equal volumes (e.g. 100 μl old medium + 100 μl diluted compound) mix well simply via pipetting. The density of DMSO is higher than the density of culture medium. Therefore, if DMSO-based solutions were applied directly to cell cultures, DMSO and compound gradients could form within the wells leading to subsequent compound precipitation, higher cytotoxicity, and/or higher compound concentrations directly above the cell layers. All of these artifacts can be avoided with our method. The highest tolerated final concentration of DMSO (usually less than 1%) depends on the cell line and must be optimized in separate experiments.

Third, the concentration of FDA, Hoechst33342 and PI together with the staining time must be optimized for each cell line studied. Since FDA itself is non-fluorescent and the action of esterase enzymes present only in living cells is necessary to hydrolyze it to fluorescein^20, 21^, short staining times may lead to insufficient signal to identify cells. If the incubation time is too long or the FDA solution is not replaced by fresh medium, cells accumulate large amounts of fluorescein leading to extremely bright fluorescence that makes it difficult to identify individual cells in the cell layer. Hoechst33342 and PI must be present at concentrations that yield effective staining within the staining time optimal for FDA.

Finally, the number of cells imaged must be high enough for each well to avoid seriously under- or overestimated signal values, which may occur solely due to the random nature of cell sampling. We have modelled the effect of sample size (number of cells/well) on the accuracy of the estimated signal by assuming that there are mono- and binucleated cells only in the cell population, which is a near-realistic assumption even after 24 h of *para*-aminoblebbistatin treatment (positive control). We further assumed that the ratio of mononucleated cells (in other words the probability of finding a mononucleated cell with random sampling) is p=0.85 and p=0.3 for negative and positive controls, respectively. This yields an expected NCR of 1.15 and 1.7, respectively. (These ratios were determined in preliminary experiments with manual cell counting.) There are exactly n+1 different possible values of the NCR for sample sizes n>2 (except for the extreme situations where all cells are mono- or binucleated). For example, if the sample size was only two cells, there would be only 3 (=2+1) different possible estimated values: 1, 1.5, and 2, for situations where 2, 1 and 0 cells are mononucleated, respectively. Note that this is always true, regardless of the true value of the NCR (e.g. 1.15 and 1.7 above). The probability (f) of finding any possible value can be calculated based on the binomial distribution:

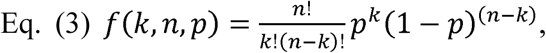

where n is the sample size, k is the number of mononucleated cells, and p is the probability of finding a mononucleated cell. Calculating for every possible value yields the probability mass function of the particular distribution. Probability mass functions calculated for several sample sizes are shown in Fig. 6A. To improve assay quality (to have a Z’ > 0.5), it is crucial to obtain “good” estimates of the NCR for every single well. These must be close enough to the true value, e.g. it must fall into the range of true value ± 5%. Since the nature of sampling is probabilistic, the sample size needs to be high enough to achieve this goal. The probabilities of having “good” estimates can be calculated as the sum of the probabilities of all the possible values of the NCR falling into the above defined range (Fig. 6B). To ensure that there is less than one well on average for each 96-well plate that might be affected by random sampling issues, this probability must be above 99%, which can be achieved by sample sizes >300/well. In this work, imaging was continued until at least 600 cells were detected in each well to make sure that final cell numbers are beyond the 300 cells/well threshold. Unfortunately, this threshold represents one of the highest limitations of the method. Since it might be necessary to image a significant part of the total area of several wells, further miniaturization (e.g. using 384 well plates) may result in less reliable data, especially when cell loss occurs due to cytotoxicity.

**Figure 6.**
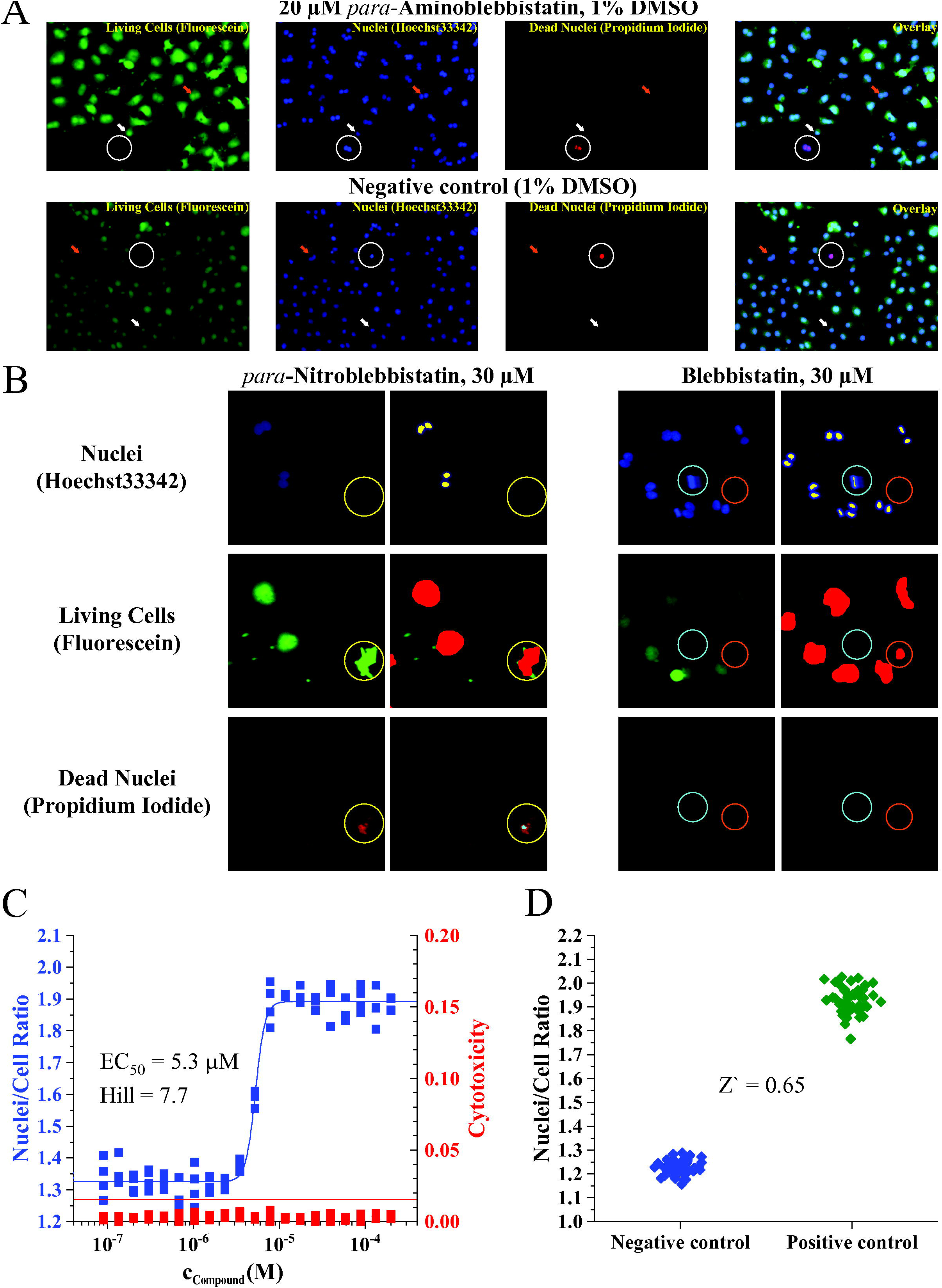
(A) Probability mass functions describing the probabilities of finding all possible NCRs depending on the sample size “n” (number of cells imaged) were calculated based on the binomial distribution. It was assumed that there are mono- and binucleated cells only and the true value of the NCR is 1.7 and 1.15 for positive and negative controls (not shown), respectively. Note that the probabilities were calculated for discrete values of the estimated NCR, as there are exactly n+1 different possible values of the NCR for sample sizes n>2 (the distributions are not continuous functions). In these calculations, the NCR was always between 1 (all cells are mononucleated) and 2 (all cells are binucleated). True values of the NCR are shown as red vertical lines. (B) The probability of finding an estimate of the NCR close to the true value can be calculated as the sum of all probabilities falling in the range of true value ±5% (defined in this work). This probability is highly dependent on the sample size for both positive (red) and negative (blue) controls. Obtaining good estimates is critical to assay reliability (Z-factor). A sample size less than ∼300 cells may result in seriously under- or overestimated parameter values solely due to the random nature of sampling.

A similar phenotypic high-throughput screen for the identification of novel cytokinesis inhibitors has already been published^29^. In that screen, fixed cells were stained by two dyes to visualize the cytoplasm and nuclei. Compared to our method, this procedure is substantially more time consuming, as it requires multiple staining and washing steps^29^. Moreover, with each of these steps, the risk of losing cells from the surface increases and, because the cells are fixed, cytotoxicity can only be estimated in separate experiments. In contrast, our method directly applies a single staining step with three dyes to living cells, thereby reducing the risk of cell loss and providing cytotoxicity information. Others have used EGFP-a-tubulin mCherry-H2B-labeled HeLa Kyoto cells to visualize the multinucleated phenotype after treating cells with blebbistatin derivatives^8, 9^. Because these cells express EGFP-a-tubulin in the cytoplasm and mCherry-H2B in nuclei, they do not require any staining before imaging. Although stain-free live cell imaging provides this advantage, one would need to introduce similar genetic modifications to any other cell lines prior to using them in screening experiments. In contrast, our strategy is easily applicable to any cell line. Interestingly, the combination of FDA, PI and Hoechst33342 dyes have recently been used to assess neuronal viability in cerebellar granule neuron cultures^30^. This indicates that the same triple-staining strategy might be useful in different areas of cell biology in the future.

In summary, the cytokinesis assay presented here is amenable to semi high-throughput screening applications. It can identify cytokinesis inhibitors regardless of their molecular target or mechanism of action. It is simple and seems to be compatible with multiple cell lines of interest, since it requires only a single staining step and relatively inexpensive, small molecule fluorescent dyes. The assay is relatively resistant to artifacts due to double fluorescent labeling and provides information about both the potency and cytotoxicity of compounds. Further miniaturization using 384 well plates may be possible depending on the characteristics of the cell line used (e.g. optimal cell density) and the cytotoxicity of the compounds tested.

